# Comparison of long-read sequencing technologies in the hybrid assembly of complex bacterial genomes

**DOI:** 10.1101/530824

**Authors:** Nicola De Maio, Liam P. Shaw, Alasdair Hubbard, Sophie George, Nick Sanderson, Jeremy Swann, Ryan Wick, Manal AbuOun, Emma Stubberfield, Sarah J. Hoosdally, Derrick W. Crook, Timothy E. A. Peto, Anna E. Sheppard, Mark J. Bailey, Daniel S. Read, Muna F. Anjum, A. Sarah Walker, Nicole Stoesser, on behalf of the REHAB consortium

**Author notes:** These authors contributed equally (Nicola De Maio, Liam P. Shaw). Corresponding authors: Nicole Stoesser.

## Abstract

Illumina sequencing allows rapid, cheap and accurate whole genome bacterial analyses, but short reads (<300 bp) do not usually enable complete genome assembly. Long read sequencing greatly assists with resolving complex bacterial genomes, particularly when combined with short-read Illumina data (hybrid assembly). However, it is not clear how different long-read sequencing methods impact on assembly accuracy. Relative automation of the assembly process is also crucial to facilitating high-throughput complete bacterial genome reconstruction, avoiding multiple bespoke filtering and data manipulation steps. In this study, we compared hybrid assemblies for 20 bacterial isolates, including two reference strains, using Illumina sequencing and long reads from either Oxford Nanopore Technologies (ONT) or from SMRT Pacific Biosciences (PacBio) sequencing platforms. We chose isolates from the Enterobacteriaceae family, as these frequently have highly plastic, repetitive genetic structures and complete genome reconstruction for these species is relevant for a precise understanding of the epidemiology of antimicrobial resistance. We *de novo* assembled genomes using the hybrid assembler Unicycler and compared different read processing strategies. Both strategies facilitate high-quality genome reconstruction. Combining ONT and Illumina reads fully resolved most genomes without additional manual steps, and at a lower consumables cost per isolate in our setting. Automated hybrid assembly is a powerful tool for complete and accurate bacterial genome assembly.

**IMPACT STATEMENT:** Illumina short-read sequencing is frequently used for tasks in bacterial genomics, such as assessing which species are present within samples, checking if specific genes of interest are present within individual isolates, and reconstructing the evolutionary relationships between strains. However, while short-read sequencing can reveal significant detail about the genomic *content* of bacterial isolates, it is often insufficient for assessing genomic *structure*: how different genes are arranged within genomes, and particularly which genes are on plasmids – potentially highly mobile components of the genome frequently carrying antimicrobial resistance elements. This is because Illumina short reads are typically too short to span repetitive structures in the genome, making it impossible to accurately reconstruct these repetitive regions. One solution is to complement Illumina short reads with long reads generated with SMRT Pacific Biosciences (PacBio) or Oxford Nanopore Technologies (ONT) sequencing platforms. Using this approach, called ‘hybrid assembly’, we show that we can automatically fully reconstruct complex bacterial genomes of Enterobacteriaceae isolates in the majority of cases (best-performing method: 17/20 isolates). In particular, by comparing different methods we find that using the assembler Unicycler with Illumina and ONT reads represents a low-cost, high-quality approach for reconstructing bacterial genomes using publicly available software.

**DATA SUMMARY:** Raw sequencing data and assemblies have been deposited in NCBI under BioProject Accession PRJNA422511 (https://www.ncbi.nlm.nih.gov/bioproject/PRJNA422511). We confirm all supporting data, code and protocols have been provided within the article or through supplementary data files.

## INTRODUCTION

The rapid development of microbial genome sequencing methods over the last decade has revolutionized infectious disease epidemiology, and whole genome sequencing has become the standard for many molecular typing applications in research and public health (1–4). Much of this evolution has been driven by the development of high-throughput, low-cost, second generation (short-read) sequencing methods, such as Illumina’s HiSeq and MiSeq platforms, which produce millions of low-error (0.1%) paired-end reads, generally 100-300bp in length. As such, Illumina sequencing has become the most widely used sequencing technology for microbial genomics. Multiple read processing algorithms now exist, typically enabling variant detection following mapping to a reference genome to assess genetic relatedness (e.g. for outbreak investigation or population genetic studies), or *de novo* assembly to facilitate the identification of important loci in the accessory genome, such as antimicrobial resistance genes (e.g. for epidemiological studies of resistance gene prevalence or for susceptibility prediction).

However, it has become clear that short-read sequencing has significant limitations depending on the bacterial species and/or epidemiological question. These limitations largely arise from the inability to fully reconstruct genomic structures of interest from short reads, including those on chromosomes and mobile genetic elements such as plasmids (5). An example where this genomic structure is highly relevant is the study of antimicrobial resistance (AMR) gene transmission and evolution in species of Enterobacteriaceae, which have emerged as a major clinical problem in the last decade (6). Short-read data from these species do not successfully facilitate assembly of the repetitive structures that extend beyond the maximum read length generated, including structures such as resistance gene cassettes, insertion sequences and transposons that are of crucial biological relevance to understanding the dissemination of key antimicrobial resistance genes.

The most widely used single molecule, long-read sequencing platforms, currently represented by Pacific Biosciences’ (PacBio) Single Molecule Real-Time (SMRT) and Oxford Nanopore Technologies’ (ONT) MinION sequencers, are often able to overcome these limitations by generating reads with a median length of 8-10kb, and as long as 100kb (5,7,8). However, the sequencing error rates of both long-read methods are much greater than Illumina (PacBio: 11-15%, raw, less in circular consensus reads (9); ONT: 5-40% (10)). Hybrid assembly, using combined short-read and long-read sequencing datasets, has emerged as a promising approach to generating fully resolved, accurate genome assemblies. With hybrid approaches, long reads provide information regarding the structure of the genome, specifically in plasmids, and short reads facilitate detailed assembly at local scales, and can be used to correct errors in long reads (11–13). The hybrid assembly tool Unicycler has been shown to outperform other hybrid assemblers in generating fully closed genomes (12).

We are not aware of any previously published direct comparisons of hybrid bacterial assemblies generated using long-read sequencing methods, yet the selection of a long-read sequencing approach has important cost, throughput and logistical implications. Currently, the two dominant long-read technologies are ONT and PacBio. The ONT MinION is a highly portable platform that has been deployed in several molecular laboratories, including those in low-income settings (14). Reported data yields of 10-30Gb and indexed barcoding now enable multiplexing of up to 12 bacterial isolates on a run (13). In contrast, the PacBio platform is non-portable but has been around longer, making it the most widely used for generating reference-grade bacterial assemblies to date (by way of example: as of 21^st^ January 2019, NCBI Assembly contains 201 *E. coli* assemblies generated with PacBio vs. 3 generated with MinION).

Here we compared different approaches for hybrid bacterial genome assembly, using ONT MinION, PacBio and Illumina HiSeq data generated from the same DNA extracts. We selected 20 bacterial isolates from four genera of the Enterobacteriaceae family of bacteria (*Escherichia, Klebsiella, Citrobacter* and *Enterobacter*) including two reference strains. These genera typically have large bacterial genomes between 5-6.5Mb with diverse sets of plasmids (15). We compared the advantages and disadvantages of ONT+Illumina versus PacBio+Illumina hybrid assembly, including the need for additional manual processing steps. We also investigated different strategies to optimize hybrid assembly using Unicycler for both long-read approaches.

## METHODS

### Bacterial isolates, DNA extraction and Illumina sequencing

For sequencing, we selected and sub-cultured 20 isolates across the four genera of interest from stocks of pure culture, stored in nutrient broth with 10% glycerol at −80°C. Sub-cultures were undertaken aerobically on Columbia blood agar at 37°C overnight. We chose two reference strains, *Escherichia coli* CFT073, and *Klebsiella pneumoniae* MGH78578, and 18 isolates that were part of a study investigating antimicrobial resistance in diverse Enterobacteriaceae from farm animals and environmental specimens (the REHAB study http://modmedmicro.nsms.ox.ac.uk/rehab; details of isolates in Table S1). These comprised *E. coli* (n=4), *K. pneumoniae* (n=2), *K. oxytoca* (n=2), *Citrobacter freundii* (n=2), *C. braakii* (n=2), *C. gillenii* (n=1), *Enterobacter cloacae* (n=3), *E. kobei* (n=2). We chose to investigate Enterobacteriaceae isolates as these bacteria are genetically complex: their genomes commonly contain multiple plasmids and repeat structures of varying size, making them difficult to assemble using other methods (5).

DNA was extracted from sub-cultured isolates using the Qiagen Genomic tip 100/G kit (Qiagen, Valencia, CA, USA) to facilitate long-fragment extraction. Quality and fragment length distributions were assessed using the Qubit fluorometer (ThermoFisher Scientific, Waltham, MA, USA) and TapeStation (Agilent, Santa Clara, CA, USA).

All DNA extracts were sequenced using the Illumina HiSeq 4000, generating 150bp paired-end reads. Libraries were constructed using the NEBNext Ultra DNA Sample Prep Master Mix Kit (NEB, Ipswich, MA, USA) with minor modifications and a custom automated protocol on a Biomek FX (Beckman Coulter, Brea, CA, USA). Ligation of adapters was performed using Illumina Multiplex Adapters, and ligated libraries were size-selected using Agencourt Ampure magnetic beads (Beckman Coulter, Brea, CA, USA). Each library was PCR-enriched with custom primers (index primer plus dual index PCR primer (16)). Enrichment and adapter extension of each preparation was obtained using 9μl of size-selected library in a 50μl PCR reaction. Reactions were then purified with Agencourt Ampure XP beads (Beckman Coulter, Brea, CA, USA) on a Biomek NXp after 10 cycles of amplification (as per Illumina recommendations). Final size distributions of libraries were determined using a TapeStation system as above and quantified by Qubit fluorometry.

### ONT library preparation and sequencing

ONT sequencing libraries were prepared by multiplexing DNA extracts from four isolates per flowcell using the SQK-LSK108 and EXP-NBD103 kits according to the manufacturer’s protocol with the following amendments: input DNA (1.5μg) was not fragmented, 2ml Eppendorf DNA LoBind tubes (Eppendorf, Hamburg, Germany) were used, all reactions were purified using 0.4x Agencourt AMPure XP beads, incubation time with Agencourt AMPure XP beads was doubled, elution volumes were reduced to the minimum required for the subsequent step, and elution was heated to 37°C. Libraries were loaded onto flow cell versions FLO-MIN106 R9.4 SpotON and sequenced for 48 hours.

### PacBio library preparation and sequencing

DNA extracts were initially sheared to an average length of 15kb using g-tubes, as specified by the manufacturer (Covaris, Woburn, MA, USA). Sheared DNA was used in SMRTbell library preparation, as recommended by the manufacturer. Quantity and quality of the SMRTbell libraries were evaluated using the High Sensitivity dsDNA kit and Qubit fluorometer and DNA 12000 kit on the 2100 Bioanalyzer (Agilent, Santa Clara, CA, USA). To obtain the longest possible SMRTbell libraries for sequencing (as recommended by the manufacturer), a further size selection step was performed using the PippinHT pulsed-field gel electrophoresis system (Sage Science, Beverley, MA, USA), enriching for the SMRTbell libraries >15kb for loading onto the instrument. Sequencing primer and P6 polymerase were annealed and bound to the SMRTbell libraries, and each library was sequenced using a single SMRT cell on the PacBio RSII sequencing system with 240-minute movies.

### Read preparation and hybrid assembly

ONT fast5 read files were base-called with Albacore (v2.0.2, https://github.com/JGI-Bioinformatics/albacore), with barcode demultiplexing and fastq output. Adapter sequences were trimmed with Porechop (v0.2.2, https://github.com/rrwick/Porechop). Read quality was calculated with nanostat (v0.22, https://github.com/wdecoster/nanostat) (17).

Long reads from both ONT and PacBio were prepared using four alternative strategies:

- **Basic**: no filtering or correction of reads (i.e. all long reads available used for assembly).
- **Corrected**: Long reads were error-corrected and subsampled (preferentially selecting longest reads) to 30-40x coverage using Canu (v1.5, https://github.com/marbl/canu) (7) with default options.
- **Filtered**: long reads were filtered using Filtlong (v0.1.1, https://github.com/rrwick/Filtlong) by using Illumina reads as an external reference for read quality and either removing 10% of the worst reads or by retaining 500Mbp in total, whichever resulted in fewer reads. We also removed reads shorter than 1kb and used the --trim and --split 250 options.
- **Subsampled**: we randomly subsampled long reads to leave approximately 600Mbp (corresponding to a long read coverage around 100x).

Hybrid assembly for each of the two long-read sequencing technologies and for each of the four read processing strategies (for a total of 8 hybrid assemblies per isolate) was performed using Unicycler (v0.4.0) (12) with default options.

We used Bandage (v0.8.1) (18) to visualize assemblies, and the Interactive Genome Viewer (IGV, v2.4.3) (19) to visualize discrepancies in assemblies produced by the different methods.

To simulate the effect of additional multiplexing on ONT data and assembly (with current kits allowing for up to 12 isolates to be indexed), we randomly subsampled half or one third of the ONT reads from each isolate and repeated the assembly as in the “Basic” strategy above.

### Assembly comparison

We used multiple strategies to compare the features of different hybrid assemblies of the same DNA extract. Firstly, we considered the completeness of an assembly i.e. specifically whether all contigs reconstructed by Unicycler were identified as circular structures. Circular structures typically represent completely assembled bacterial chromosomes and plasmids; circular structures from different assemblies in our 20 isolates tended to agree in the majority of cases (Table 1) and agreed with the structures of reference genomes for the two reference strains (CFT073 and MH78578). We therefore also used the number of circular contigs in an assembly as a measure of its completeness.

**Table 1.** Summary of all assemblies in terms of circularised contigs. Different rows refer to different isolates. “n of m” means that *n* contigs were circular in the assembly out of *m* total contigs. When *n* and *m* are identical, it means that the assembly was considered complete, and these cases are shaded in green. “Basic”, “Corrected”, “Filtered” and “Subsampled” refer to the strategies of long read preparation (see Methods). “NA” refers to cases where the assembly pipeline repeatedly failed. The true number of circular structures was estimated by inspection.

|  | ONT (MinION) |  |  |  | PacBio (RSII System) |  |  |  | True circular structures (estimated) |
| --- | --- | --- | --- | --- | --- | --- | --- | --- | --- |
| Isolate | Basic | Corrected | Filtered | Subsampled | Basic | Corrected | Filtered | Subsampled |  |
| CFT073 (reference) | 1 of 1 | 1 of 1 | 0 of 9 | 1 of 1 | 0 of 9 | 0 of 9 | 0 of 9 | 0 of 9 | 1 |
| MGH78578 (reference) | 6 of 6 | 4 of 7 | NA | 6 of 6 | 4 of 7 | 2 of 22 | 2 of 22 | 2 of 22 | 6 |
| RBHSTW-00029 | 3 of 9 | 3 of 9 | 3 of 9 | 3 of 9 | 3 of 9 | 3 of 9 | 3 of 9 | 3 of 9 | 4 |
| RBHSTW-00053 | 6 of 6 | 6 of 6 | 6 of 6 | 6 of 6 | 6 of 6 | 6 of 6 | 6 of 6 | 6 of 6 | 6 |
| RBHSTW-00059 | 5 of 5 | 5 of 5 | 5 of 5 | 5 of 5 | 5 of 5 | 5 of 5 | 5 of 5 | 5 of 5 | 5 |
| RBHSTW-00122 | 4 of 4 | 4 of 4 | 4 of 4 | 4 of 4 | 4 of 4 | 4 of 4 | 4 of 4 | 4 of 4 | 4 |
| RBHSTW-00123 | 7 of 7 | NA | 7 of 7 | 7 of 7 | 5 of 8 | 4 of 18 | 4 of 18 | 4 of 18 | 7 |
| RBHSTW-00127 | 5 of 5 | 5 of 5 | 5 of 5 | 5 of 5 | 5 of 5 | 5 of 5 | 5 of 5 | 5 of 5 | 5 |
| RBHSTW-00128 | 4 of 4 | 4 of 4 | 4 of 4 | 4 of 4 | 4 of 4 | 3 of 6 | 3 of 6 | 3 of 6 | 4 |
| RBHSTW-00131 | 4 of 4 | 2 of 7 | 4 of 4 | 4 of 4 | 3 of 15 | 4 of 5 | 3 of 15 | 2 of 15 | 4 |
| RBHSTW-00142 | 7 of 7 | 5 of 25 | 7 of 7 | 7 of 7 | 4 of 24 | 4 of 58 | 4 of 24 | 4 of 27 | 7 |
| RBHSTW-00167 | 9 of 9 | 5 of 15 | 10 of 10 | 9 of 9 | 4 of 34 | 3 of 60 | 3 of 60 | 3 of 60 | 9 |
| RBHSTW-00189 | 6 of 6 | 6 of 6 | 5 of 6 | 6 of 6 | 5 of 29 | 5 of 28 | 5 of 29 | 5 of 30 | 6 |
| RBHSTW-00277 | 2 of 2 | 2 of 2 | 1 of 8 | 2 of 2 | 1 of 8 | 1 of 8 | 1 of 8 | 1 of 8 | 2 |
| RBHSTW-00309 | 4 of 5 | 5 of 5 | 5 of 5 | 4 of 5 | 5 of 5 | 4 of 5 | 5 of 5 | 5 of 5 | 5 |
| RBHSTW-00340 | 3 of 11 | 3 of 11 | 4 of 4 | 4 of 4 | 2 of 25 | 2 of 25 | 2 of 24 | 2 of 25 | 4 |
| RBHSTW-00350 | 2 of 2 | 2 of 2 | 2 of 3 | 2 of 2 | 2 of 2 | 2 of 2 | 2 of 2 | 2 of 2 | 2 |
| RHB10-C07 | 1 of 1 | 1 of 1 | 1 of 1 | 1 of 1 | 1 of 1 | 1 of 1 | 1 of 17 | 1 of 1 | 1 |
| RHB11-C04 | 3 of 3 | 3 of 3 | 3 of 3 | 3 of 3 | 3 of 3 | 3 of 3 | 3 of 3 | 3 of 3 | 3 |
| RHB14-C01 | 1 of 12 | 1 of 12 | 1 of 15 | 1 of 12 | 1 of 15 | 1 of 15 | 1 of 15 | 1 of 15 | 2 |
| Total contigs | 109 | 130 | 115 | 102 | 218 | 294 | 276 | 265 | 87 |

|  | ONT (MinION) |  |  |  | PacBio (RSII System) |  |  |  |
| --- | --- | --- | --- | --- | --- | --- | --- | --- |
| Total circularised contigs (% over total estimated circular structures from Bandage: n=87 for all isolates) | 83 (95%) | 67 (84%) | 77 (95%) | 84 (97%) | 67 (77%) | 62 (71%) | 62 (71%) | 61 (70%) |
| Total circularised contigs for reference strains (i.e. structures known, total n=1 [*E. coli*] + 6 [*K. pneumoniae*]) | 7 (100%) | 5 (71%) | 0 (0%) | 7 (100%) | 5 (71%) | 2 (29%) | 2 (29%) | 2 (29%) |
| Total isolates with all contigs circularised (% isolates) | 16 (80%) | 12 (60%) | 13 (65%) | 17 (85%) | 9 (45%) | 7 (35%) | 7 (35%) | 8 (40%) |

A common error associated with long-read-based assemblies is indel errors, which can artificially shorten proteins by introducing premature stop codons or frameshift errors (20). To check this possibility we annotated genomes with Prokka (v1.13.3, https://github.com/tseemann/prokka) (21) then aligned all proteins to the full UniProt TrEMBL database (November 15^th^ 2018) using DIAMOND (v0.9.22, https://github.com/bbuchfink/diamond) (22) and compared the length of each protein to its top hit. We compared proteins in assemblies for the same sample with Roary (v3.12.0, https://sanger-pathogens.github.io/Roary) (23).

We additionally compared different assemblies of the same extract using:

- ALE (24), which assesses the quality of different assemblies using a likelihood-based score of how well Illumina reads map to each assembly. ALE was run with default parameters; Illumina reads were mapped to references using Bowtie2 (v2.3.3) (25).
- DNAdiff (as part of MUMMER v3.23) (26), which compares assemblies of the same strain to detect differences such as SNPs and indels. DNAdiff was run with default parameters on the fasta assembly files.
- REAPR (v1.0.18) (27), which (similarly to ALE) evaluates the accuracy of assemblies using information from short read mapping to the assembly. REAPR was run using the options “facheck”, “smaltmap” and “pipeline” with default parameters.
- Minimap2 (v2017-09-21) (28) was used to map long reads to the hybrid assemblies, and the mappings were evaluated to compare assembly quality and long read features (identity and length) using scripts from the Filtlong package. We considered the average identity for each base, and if there were multiple alignments at a base, we used the one with the best score. We aligned PacBio and ONT reads to the hybrid assemblies obtained either from all PacBio reads or from all ONT reads. Read alignments were classified as: “good” if they had at least one alignment covering 97% of the read, as a putative “chimera” if they had multiple inconsistent alignments represented by at least 10% of the read length and ≥70% nucleotide identity, and “other” if they did not fall into either of the two previous categories.

## RESULTS

### Sequencing data quality

For Illumina data, a median of 2,457,945 (interquartile range [IQR]: 2,073,342-2,662,727) paired reads was generated for each isolate, with a median insert size of 363 bp (351-369). The %GC content per isolate varied, as expected, by genus (median 53%, range: 50-57%), but was consistent with the expected %GC content for each isolate based on its species (Table S1).

The PacBio SMRT sequencing data resulted in a median of 160,740 (IQR: 153,196-169,240) sub-reads with median sub-read length of 11,050 bp (IQR: 10,570-11,209 bp) per isolate. Each isolate was sequenced using one SMRT cell on the RSII sequencing system, generating a median of 1.32Gb (IQR: 1.25-1.36) of data per isolate, with isolates being run in batches of 8 (Figure S1, Table S1). For the ONT data, a median of 102,875 reads (IQR: 70,508-143,745 reads) were generated for each isolate, with a median phred score of 11.8 (IQR: 11.4-12.3). ONT reads had a median length of 14,212 bp (IQR: 13,369-16,267 bp). A median of 13.8Gb (IQR: 10.8-14.7Gb) of data was generated per run, resulting in a median of 3.45Gb per isolate (four isolates multiplexed per run) (Figure S1, Table S1). After hybrid assembly, the mean percentage identity and identity N50 for reads aligned against their respective assemblies were higher for ONT reads than PacBio reads (mean±s.d. read alignment identity: 86±7 vs. 78±17; Figure S3, Table S3).

### Hybrid assembly runtimes

Clearly the computing infrastructure available to any given research team will be widely variable, and assembly runtimes will therefore be different. For this experiment, where all assemblies were run with dual 8-core Intel IvyBridge 2.6GHz, 256GB 1866MHz memory, assembly times averaged between 1600-8000 minutes (~26-130 hours, Table S4), depending on long-read preparation strategy (i.e. basic, corrected, filtered, sub-sampled, as in Methods). They did not significantly vary depending on type of long-read used as input. Assemblies completed in all cases, apart from two cases (both ONT+Illumina hybrids: MGH78578 reference strain, filtered strategy; RBHSTW-00123, corrected strategy).

### PacBio vs. ONT-based hybrid assembly comparisons

Using ONT+Illumina hybrid assembly approaches, we were able to completely assemble (i.e. all contigs circularised) the majority of genomes (between 12 [60%] and 17 [85%] depending on the preparation strategy for long reads, Table 1) without any manual intervention (18 across all strategies). With PacBio+Illumina fewer assemblies were complete (between 7 [35%] and 9 [45%]). More contigs were also circularised with ONT than with PacBio (up to 84 [97%] versus 67 [77%]), and assemblies were less fragmented (a minimum of 102 total contigs across all 20 isolates for ONT vs. a minimum of 218 for PacBio).

On the basis of the minimap2/Filtlong comparisons (see Methods), most reads from both long-read platforms had “good” alignment to their respective assemblies (~103,000 reads on average for PacBio vs. ~99,000 reads for ONT, Figure S2, Table S2), with slightly more alignments classified as “chimeras” (4,502 vs. 1,074 reads) and a much larger number of alignments that were poor and classified as “other” (54,449 vs. 8,222) for PacBio compared to ONT reads (Figure S2, Table S2).

Some chromosomal regions proved hard to assemble with both PacBio and ONT, e.g. for isolates RBHSTW-00029 and RHB14-C01, but one of the noticeable differences between the two methods was the ability of ONT to resolve repeats on small plasmids (see Figure 1 and Figure S4). The DNA fragment size selection process used to optimize PacBio sequencing and recommended by the manufacturer may have contributed to this (see Methods), essentially making the assembly of small plasmids reliant on the Illumina short-read component of the dataset only. This also reduces the power of PacBio reads for resolving the genome structure when one copy of a repeated region is present on a short plasmid.

**Figure 1.**
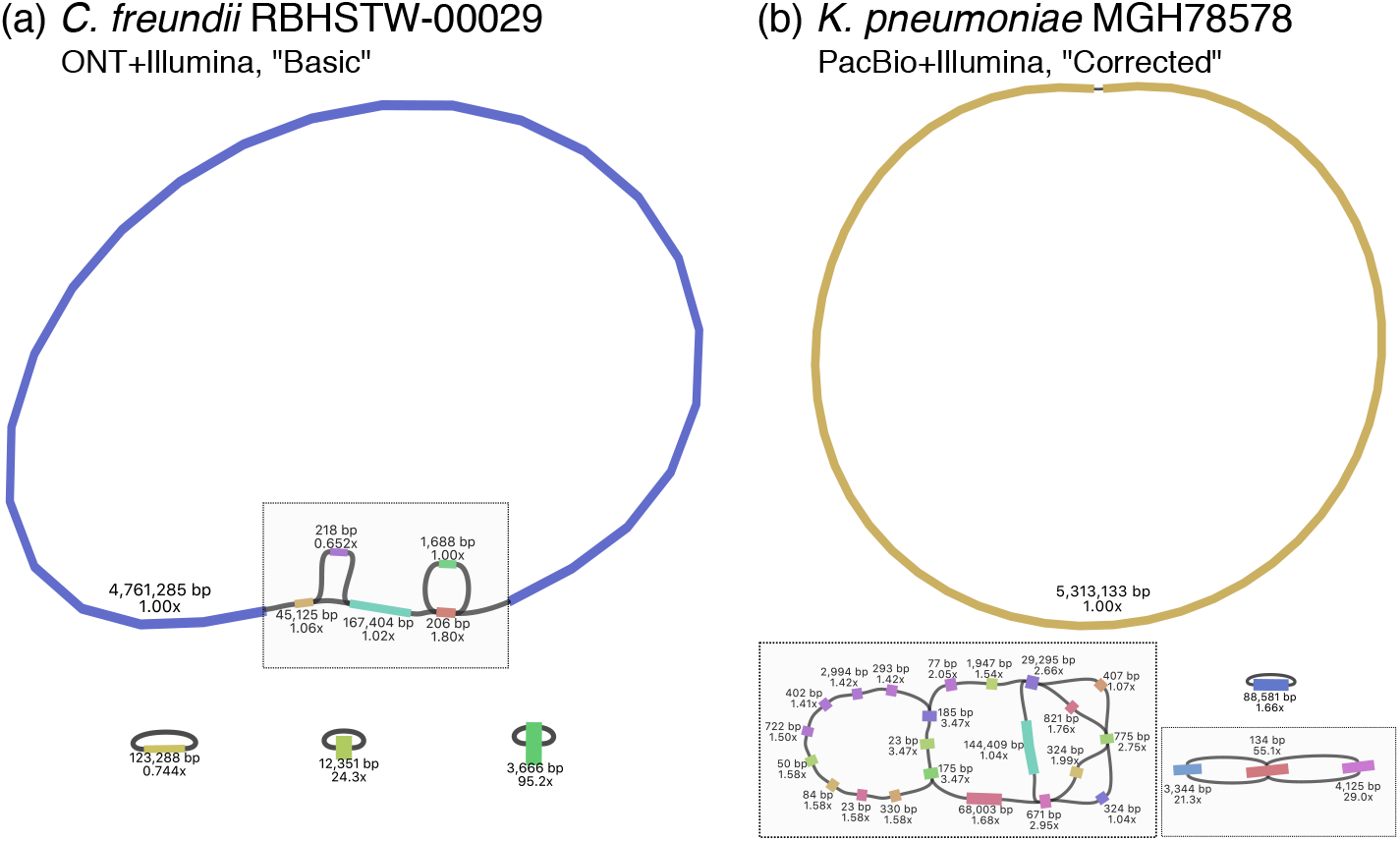
Examples of genome structure uncertainty in hybrid assemblies in a) the chromosome and b) the accessory genome. (a) An ONT+Illumina hybrid assembly for isolate RBHSTW-00029 using the “Basic” long read preparation strategy. b) A PacBio+Illumina hybrid assembly for isolate MGH78578 using the “Corrected” long read preparation strategy. Plots were obtained using Bandage, with grey boxes indicating unresolved structures. Each contig is annotated with contig length and Illumina coverage; connections between contigs represent overlaps between contig ends. The assembly for RHBSTW-00029 in a) and that of isolate RHB14-C01 (which showed a similar pattern of chromosome structure uncertainty) represented the only two datasets that could not be completely assembled with any of the attempted strategies using ONT+Illumina data. They were also not fully assembled by any PacBio+Illumina strategy, which similarly failed to completely assemble isolates RBHSTW-00189, RBHSTW-00277, RBHSTW-340 and CFT073 (Figure S4). The pattern in b) was only observed for PacBio+Illumina data, and was the reason for incomplete assemblies for isolates RBHSTW-00123, RBHSTW-00131, RBHSTW-00142, RBHSTW-00167 and MGH78578 (Figure S4).

While correcting ONT reads with Canu or filtering them with Filtlong improved assembly completeness for one isolate (RBHSTW-00309), in most cases avoiding this ONT read correction and filtration led to better results (Table 1). This might be due to correction and filtration steps removing reads in a non-uniform way across the genome, and in particular from regions that are already hard to assemble. An alternative strategy deployed to reduce the computational burden of hybrid assembly was to randomly sub-sample long reads until a certain expected coverage was reached. Table 1 shows that this strategy was preferable to read correction and filtration: it did not reduce assembly completeness but did reduce computational demand (from an average of 5640 minutes to 2020 minutes per assembly on a dual 8-core Intel IvyBridge 2.6GHz, 256GB 1866MHz memory, Table S4).

The analysis of local sequence assembly quality was inconclusive, showing inconsistent results across different methodologies (Table 2), suggesting neither approach was clearly superior to the other in this respect. However, detailed investigation of single nucleotide polymorphisms (SNPs) between ONT and PacBio-based assemblies for the reference isolates demonstrated two specific patterns of assembly differences. First, some positions (17 SNPs across the two reference isolates) appeared plausibly polymorphic in the original DNA sample and were called differently in different assembly runs (see Figure 2a). Secondly, positions within regions with extremely low Illumina coverage (see Figure 2b) could have led to assembly errors (25 SNPs across the two reference isolates), the PacBio assemblies being more affected (22 cases vs 3 for ONT).

**Figure 2.**
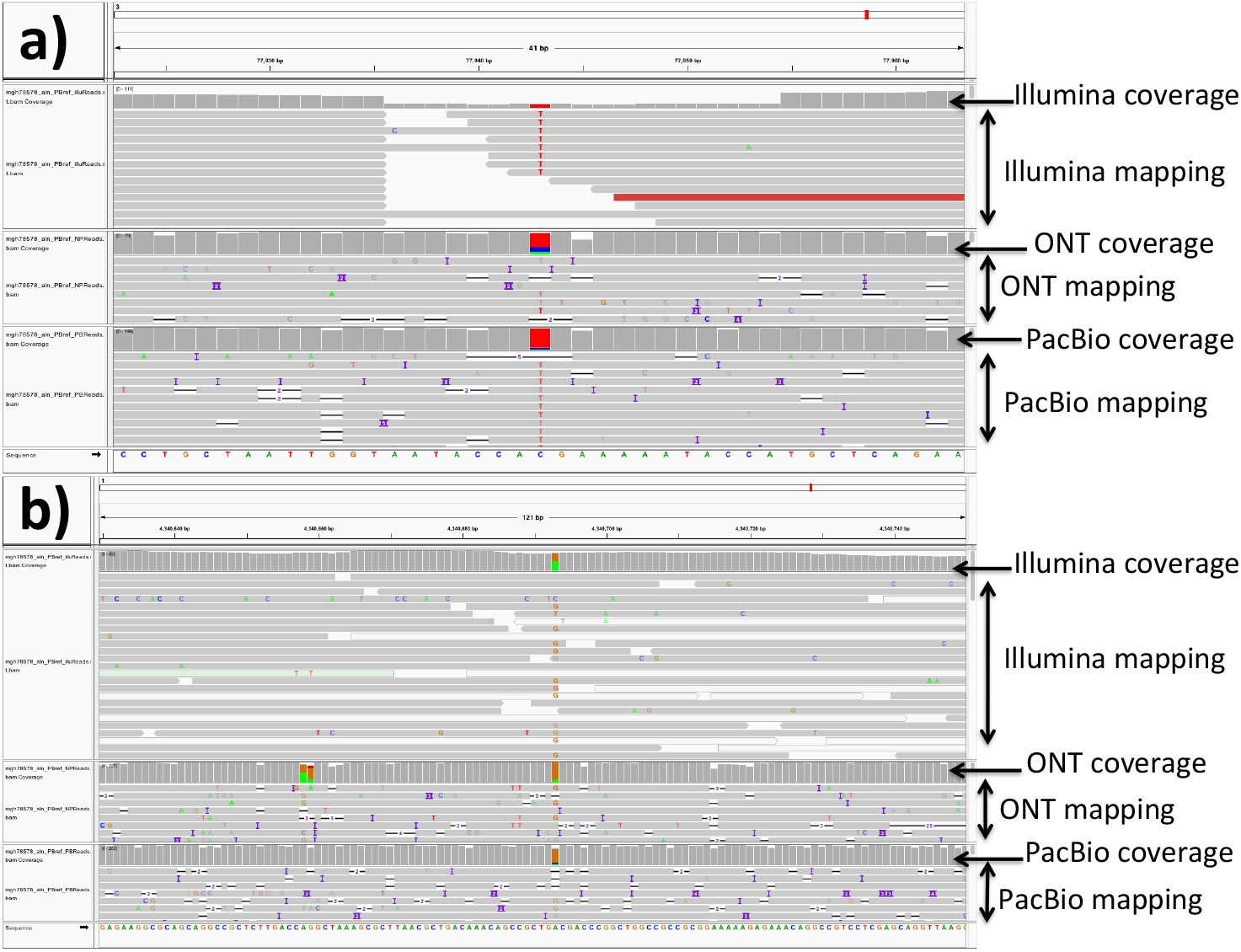
Examples of mismatches identified between the ONT-based and the PacBio-based assemblies for the two reference strains (*E. coli* CFT073 and *K. pneumoniae* MGH78578). Each sub-figure is an IGV (v2.4.3) view of part of the PacBio-based assembly, centered around a PacBio-ONT SNP, with all reads from the same isolate mapped to it. We performed this analysis for all SNPs in isolates MGH78578 and CFT073, and report examples for the two most typical patterns observed. a) SNP from MGH78578 with very low Illumina coverage, but normal PacBio and ONT coverage. Most of the Illumina reads have a different base than the one in the PacBio-assembled reference (the red T’s), suggesting perhaps an error in the PacBio assembly. A similar pattern is observed in 14 SNPs in CFT073 (with 12 due to error in the PacBio assembly), and 11 SNPs in MGH78578 (with 10 due to error in the PacBio assembly). b) SNP from MGH78578 with normal Illumina coverage; Illumina reads support both bases with similar proportions, suggesting that this could be a polymorphic site within the original DNA sample. This pattern was observed for 4 SNPs in CFT073 and for 13 SNPs in MGH78578.

**Table 2.** Comparison between PacBio and ONT-based hybrid assemblies. Comparisons are shown using ALE, DNAdiff and REAPR (see Methods). Different rows represent different isolates. All entries representing a better score for the PacBio assembly are shaded in red, those showing a better score for ONT are shaded in blue. “ALE score” is the assembly likelihood difference (calculated by ALE from the mapping of Illumina reads) between PacBio and ONT assemblies. “Unmapped reads” refers to the number of Illumina reads that ALE did not map to the corresponding assembly. “REAPR errors” refers to the assembly errors found by REAPR by mapping Illumina reads to the corresponding assembly. For each isolate, one ONT and one PacBio-based assembly with the best completion (i.e. number of circularised contigs) were chosen for comparison. DNAdiff results show the median (range) results from comparing all assemblies for an isolate across read preparation strategies i.e. 4×4=16 comparisons for each isolate. “GSNPs” / “GIndels” refer to high-confidence SNPs / indels between ONT and PacBio assemblies.

| Isolate | ALE score | PacBio unmapped reads (% total) | ONT unmapped reads (% total) | PacBio REAPR errors | ONT REAPR errors | DNAdiff GSNPs | DNAdiff GIndels |
| --- | --- | --- | --- | --- | --- | --- | --- |
| CFT073<br>(reference *E. coli*) | -17928 | 29246 (0.89%) | 29240 (0.89%) | 5 | 5 | 1 (0-1) | 0 (0-0) |
| MGH78578<br>(reference *K. pneumoniae*) | -1532602 | 41793 (1.31%) | 38371 (1.21%) | 8 | 7 | 6 (1-7) | 0 (0-1) |
| RBHSTW-00029 | 207465 | 50056 (1.85%) | 49876 (1.84%) | 3 | 3 | 0 (0-0) | 0 (0-0) |
| RBHSTW-00053 | 4727 | 50860 (1.62%) | 50861 (1.62%) | 12 | 11 | 1.5 (0-4) | 0 (0-0) |
| RBHSTW-00059 | -143627 | 37357 (1.04%) | 36251 (1.01%) | 15 | 14 | 0 (0-0) | 0 (0-0) |
| RBHSTW-00122 | 0 | 24355 (1.18%) | 24355 (1.18%) | 6 | 7 | 0 (0-0) | 0 (0-0) |
| RBHSTW-00123 | -1963188 | 56224 (1.68%) | 57074 (1.70%) | 17 | 21 | 4 (1-6) | 4.5 (2-6) |
| RBHSTW-00127 | -1145 | 34206 (0.98%) | 34206 (0.98%) | 16 | 16 | 0 (0-0) | 0 (0-0) |
| RBHSTW-00128 | 3114 | 31526 (1.06%) | 31507 (1.05%) | 6 | 8 | 2 (1-2) | 2 (1-4) |
| RBHSTW-00131 | 399368 | 25880 (0.88%) | 26271 (0.89%) | 24 | 28 | 3 (1-7) | 1 (1-3) |
| RBHSTW-00142 | -790773 | 34684 (1.23%) | 32590 (1.16%) | 12 | 12 | 3 (1-11) | 0 (0-1) |
| RBHSTW-00167 | 4083063 | 34510 (1.13%) | 76805 (2.52%) | 24 | 33 | 21 (18-47) | 1.5 (0-4) |
| RBHSTW-00189 | -158523 | 37378 (1.25%) | 37418 (1.25%) | 9 | 12 | 11.5 (7-21) | 1 (0-2) |
| RBHSTW-00277 | 18417 | 33677 (0.99%) | 33685 (0.99%) | 16 | 16 | 2 (0-2) | 0 (0-0) |
| RBHSTW-00309 | -518811 | 30704 (0.88%) | 30327 (0.87%) | 17 | 36 | 2 (0-11) | 44.5 (0-86) |
| RBHSTW-00340 | -906675 | 30802 (0.87%) | 29860 (0.84%) | 11 | 10 | 2 (0-4) | 0 (0-1) |
| RBHSTW-00350 | 21188 | 28907 (0.79%) | 28907 (0.79%) | 12 | 13 | 2 (2-4) | 5 (0-8) |
| RHB10-C07 | -23295 | 27779 (0.90%) | 27777 (0.90%) | 22 | 21 | 5 (0-17) | 0.5 (0-1) |
| RHB11-C04 | 12774 | 24879 (0.86%) | 24881 (0.86%) | 25 | 25 | 2 (0-6) | 0 (0-0) |
| RHB14-C01 | 172712 | 30478 (0.95%) | 30576 (0.95%) | 13 | 12 | 3 (0-3) | 0 (0-0) |

The proportion of proteins with a length of <90% of their top UniProt hit was low (~2-4% c.f. 3.7% for the RefSeq assembly of *E. coli* MG1655) and extremely consistent across ONT+Illumina and PacBio+Illumina assemblies (Figure S5), suggesting that indels were not a significant problem in the assemblies. There was very close agreement between methods (median discrepancy < 5 proteins), although there were a greater number of cases where more proteins were found in the ONT+Illumina assemblies (Figure S6). Proteins found uniquely in an assembly tended to be found on a contig that was fragmented in the comparison assembly (e.g. the third plasmid in the ONT-based assembly for RBHSTW-00167 was fragmented in the comparison PacBio-based assembly, and was the location of 11 proteins unique to the ONT-based assembly), highlighting that the degree of contig fragmentation in an assembly can affect conclusions about gene presence beyond just the inability to resolve genomic structures (Table S5, Figure S4).

Comparing *de novo* assemblies and reference genomes for the two reference strains (CFT073 and MGH78578) we found that the hybrid assemblies from ONT and PacBio reads were more similar to each other (e.g. 18 SNPs and 0 indels for CFT073 and 24 SNPs and 13 indels for MGH78578) than to the available reference genome sequences (156-365 SNPs and 47-439 indels vs. the references, Table S6), possibly due to: (i) strain evolution in storage and sub-culture since the reference strains were sequenced; (ii) errors in the original reference sequences; and/or (iii) consistent errors in the hybrid assemblies.

Lastly, we investigated the effects of further ONT multiplexing by simulating datasets with 8 and 12 barcodes respectively (see Methods). Halving the available reads (equivalent to 8 barcodes) had no negative effect on the assemblies (Table S7). Using a third (equivalent to 12 barcodes) slightly increased the fragmentation of the assemblies overall (one fewer completed assembly and nine additional non-circular contigs). However, these results were not uniform: two assemblies gained an extra circular contig (RBHSTW-00309 and RBHSTW-00340) with this downsampling.

### DNA preparation and sequencing costs

Beyond considerations of assembly accuracy, an important and realistic consideration when choosing a sequencing approach is cost. While we do not attempt to calculate estimates that will apply across different labs and settings, we can report our consumables costs per isolate (i.e. exclusive of other potential costs, such as labour/infrastructure [laboratory and computational]) in case it is helpful for informing others. The cost of bacterial culture and DNA extraction was approximately £12 per isolate, resulting in sufficient DNA for all three sequencing methods to be performed in parallel on a single extract. Cost for Illumina library preparation and sequencing (see Methods) was ~£41 per isolate. ONT MinION sequencing (library preparation and run) was performed by multiplexing 4 isolates per run, resulting in costs of approximately £130 per isolate; however, it is possible to multiplex up to 12 isolates per run at correspondingly lower coverage (13), resulting in costs of ~£44/isolate. At the time we performed these experiments (late 2017), the PacBio sequencing was done using one isolate per library per SMRTcell on the RSII system, with PacBio sequencing costs of more than £280 per isolate. However, at the time of manuscript preparation, microbial sequencing had been transferred to the higher throughput PacBio Sequel system, on which multiple isolates can be multiplexed per SMRTcell 1M. Assuming ownership of a Sequel system, the updated cost for PacBio sequencing, including DNA fragmentation, SMRTbell preparation, size selection on the BluePippin system (Sage Science) and sequencing, is £190 per isolate when multiplexing 8 isolates. If less coverage is needed or smaller genomes are to be examined, one could multiplex up to 16 isolates per SMRTcell 1M at a cost of £152 per isolate.

To summarise, in the optimal scenario for each technology in our setting, our total predicted consumables costs range from £97-183 for generating an ONT+Illumina hybrid assembly (multiplexing 4 versus 12 isolates) to £205-255 for generating a PacBio+Illumina hybrid assembly on the PacBio Sequel system (multiplexing 8 versus 16 isolates). Costs using the PacBio RSII system (i.e. >£320) to generate PacBio+Illumina hybrid assemblies would be substantially higher than those for generating an ONT+Illumina hybrid assembly. We stress that these costs are estimates only, and specifically do not include infrastructural and staffing costs.

## DISCUSSION

Combining short read Illumina sequencing with different long read sequencing technologies and using Unicycler, a publicly available and widely-used hybrid assembly tool, we found that ONT+Illumina hybrid assembly generally facilitates the complete assembly of complex bacterial genomes without additional manual steps. Our data thus support ONT+Illumina sequencing as a non-inferior bacterial genome hybrid assembly approach compared with PacBio+Illumina, leading to more complete assemblies, and to significantly lower costs per isolate if multiplexed.

We also investigated the impact of different long-read processing strategies on assembly quality and found that different strategies can result in more complete assemblies. We showed that quality-based filtration and correction of long reads can apparently paradoxically result in worse performance than just using unfiltered and uncorrected reads. There is no obvious explanation for this; although we speculate that preferential removal of long reads from hard-to-sequence regions might be a contributing factor, we have been unable to establish if this is the case. We propose a different strategy to reduce the computational burden of hybrid assembly without affecting the final outcome, namely randomly sub-sampling long reads down to a desired level of coverage. We demonstrated that this strategy generally results in better assemblies for ONT sequencing data.

We did however identify some recurrent patterns of local hybrid misassembly that could be systematically addressed in the future. One of these is the presence of polymorphisms in the DNA extract. These may represent genuine minor variants present in the isolate (although it is difficult to establish with certainty), but the salient fact here is that current bacterial assembly methods assume that no position is polymorphic which can lead to an imperfect representation of the genomic content where this is not the case. We advocate for the inclusion or awareness of polymorphisms within assembly polishing methods e.g. Pilon (29).

The other problem we identified is that regions with very low Illumina coverage tend to be enriched with small assembly errors. This problem could similarly be addressed in the future with hybrid assembly polishing methods, which would supplement Illumina-based polishing with long read-based polishing in regions with low Illumina coverage.

There were several limitations to our study. Firstly, we included only two reference strains, and our analyses suggest that the “true” sequences for these had diverged from the publicly available reference sequences. This divergence could arise from multiple sources: true biological variation after years of storage and/or sub-culture (a known possibility that has been previously observed for bacterial reference strains e.g. in archived cultures of *Salmonella enterica* serovar Typhimurium LT2 (30)), errors in the original reference sequences (first published in 2002 for CFT073, 2007 for MGH78578), or possible errors in our hybrid assemblies. Thus, making comparisons for any given approach even in the case where a reference is available is difficult in the absence of a clear gold standard. Of note, we tried to minimize biological variability introduced in culture by sequencing the same DNA extract across different platforms. For 18 isolates the “true” underlying sequence was unknown, which is common for highly plastic Enterobacteriaceae genomes. There is no consensus on how best to evaluate assemblies and assembly quality when a reference is not available. We therefore used several approaches, and these were not always consistent with each other.

Assemblies can sometimes be further improved after an initial evaluation using “manual completion” (see https://github.com/rrwick/Unicycler/wiki/Tips-for-finishing-genomes). However, we did not investigate manual completion for our hybrid assemblies because, in general, it is hard to replicate, has not been benchmarked and validated, is more easily biased, and is not feasible for processing large numbers of isolates. We did not identify any published, publicly available tools developed to specifically handle PacBio+Illumina hybrid assembly, although some research groups may have implemented and validated these in-house. Finally, we did not investigate the effect of different basecallers. The evolution of both technologies and post-sequencing processing of data generated by both ONT and PacBio platforms is rapid, and recent advances have been made e.g. in basecalling with the switch from Albacore to Guppy for ONT data. Our assumption is that such advances which improve read quality and basecalling will improve assembly quality, but we have not carried out specific comparisons.

In conclusion, we have demonstrated that reference-grade, complete hybrid assemblies can be effectively generated for complex bacterial genomes including multiple plasmids using ONT platforms in combination with Illumina data. Given the average yields that can be generated with these devices, it should be feasible to comfortably multiplex eight Enterobacteriaceae isolates per ONT flowcell. At current listed cost prices, this effectively represents a cost of ~£100/hybrid assembly (all laboratory and sequencing consumables costs [includes Illumina and Nanopore]).

## AUTHOR STATEMENTS

### Authors and contributors

Conceptualisation: NdM, ASW, TEAP, DWC, NSt; Methodology: NdM, LPS, RW, NSt; Software: NdM, LPS, RW, AS, NSa, JS; Formal analysis: NdM, LPS, NSt; Investigation: AH, SG, MAb, ES; Resources: MA, DR, DWC, ASW, TEAP, SJH, NSt; Data curation: MAb, ES, NdM, LPS, NSt; Writing - original draft preparation: NdM, LPS, SG, ASW, NSt; Writing - review and editing: All authors; Visualisation: NdM, LPS, NSt; Supervision: MA, DR, DWC, NSt; Project administration: SJH, MA, DR, NSt; Funding: MA, DR, DWC, NSt.

### Conflicts of interest

The authors have no conflicts of interest to declare.

### Funding information

This work was funded by the Antimicrobial Resistance Cross-council Initiative supported by the seven research councils [NE/N019989/1].

Crook, George, Peto, Sheppard, Walker are affiliated to the National Institute for Health Research Health Protection Research Unit (NIHR HPRU) in Healthcare Associated Infections and Antimicrobial Resistance at University of Oxford in partnership with Public Health England (PHE) [grant HPRU-2012-10041]. The views expressed are those of the author(s) and not necessarily those of the NHS, the NIHR, the Department of Health or Public Health England.

This work is supported by the NIHR Oxford Biomedical Research Centre.

## Acknowledgements

The REHAB consortium is represented by (bracketed individuals also included in main author list): (Abuoun M), (Anjum M), Bailey M, Brett H, Bowes M, Chau K, (Crook DW), (de Maio N), Duggett N, Gilson D, Gweon HS, (Hubbard A), (Hoosdally S), Kavanaugh J, Jones H, (Peto TEA), (Read DS), Sebra R, (Shaw LP), (Sheppard AE), Smith R, (Stubberfield E), (Swann J), (Walker AS), Woodford N.

We would like to acknowledge the support of Sebra R and Smith M in the Department of Genetics and Genomic Sciences at the Icahn School of Medical Sciences at Mt Sinai (New York, USA), who participated in the experimental design of the study and performed the long-read PacBio sequencing.

We would also like to acknowledge the support of Hartwell O and Platt J (Oxford Nanopore Technologies, Oxford, UK); however, there was no input from ONT in the design, experimental work, bioinformatics, or analyses performed in this study.

## ABBREVIATIONS

ONT: :Oxford Nanopore Technologies
PacBio: :Pacific Biosciences
SNP: :single nucleotide polymorphism
AMR: :antimicrobial resistance

## SUPPLEMENTARY FIGURES AND TABLES

**Figure S1. Read counts and read length distributions for ONT and PacBio outputs**.

**Figure S2. Summary of read-to-assembly alignments**. All assemblies considered were obtained using all reads of the given type. Reads are classified as “good” if they have at least one mapping covering 97% of the read. They are classified as a putative “chimera” if they have multiple inconsistent alignments with at least 10% of read length and 70% identity. Complete statistics from minimap2/Filtlong outputs are in Table S2.

**Figure S3. Mean percent identities and identity N50 values of ONT/PacBio reads aligned to the hybrid assemblies**. We considered the average identity for each base, and if there were multiple alignments at a base, we used the one with the best score. We aligned PacBio reads to the hybrid assembly obtained from all PacBio reads. We aligned ONT reads to the hybrid assembly obtained from all ONT reads. Identity N50 represents the percent identity for which half of the total bases are in reads with this identity value or higher. Complete statistics are in Table S3.

**Figure S4. Bandage plots for hybrid assemblies**. Each square represents one genome assembly. Shown are the ONT+Illumina (left) and PacBio+Illumina (right) assemblies for each isolate (4 columns of 5 isolates). All assembly plots are for the globally optimal long read preparation strategy for each sequencing approach i.e. “Subsampled” for ONT+Illumina and “Basic” for PacBio+Illumina (see Methods). Sequential colours for plasmids are for identical structures within isolates, but not between.

**Figure S5. Percentage of proteins with a length <90% of top UniProt hit**. Proteins in assemblies were annotated with Prokka then blasted with DIAMOND against the full UniProt database (see Methods). The proportion of proteins with a length <90% of their top UniProt hit gives a simple test for artificially shortened proteins due to indel errors in assembly. The black dashed line indicates the percentage in an existing high-quality reference genome for *E. coli* MG1655 (157 proteins out of 4240; RefSeq GCF_000005845.2). Absolute numbers were all <250; shown here is the value as a percentage of the maximum number of proteins observed in any assembly for the sample to allow comparison between different genome sizes.

**Figure S6. Comparison of discrepancy in total Prokka annotated regions across all assemblies**. The discrepancy is the number of annotated regions in the ONT+Illumina assembly minus the number of annotated regions in the PacBio+Illumina assembly. All 4×4=16 comparisons of read preparation strategies are shown.

**Table S1. Summary of sequenced isolates, DNA inputs and raw sequencing metrics**.

Statistics in this table refer to raw (i.e. unfiltered) sequencing data. ONT read statistics were generated with nanostat (v0.22).

**Table S2. Classification of long reads from PacBio and ONT**. “PB” indicates PacBio. “PB2ONT” represents PacBio reads mapped to the ONT hybrid assembly, and so on. All assemblies considered were obtained using all reads of the given type. We show the number of reads falling in different categories according to how they map to the assemblies. Reads are classified as “Good” if they have at least one mapping covering 97% of the read. They are classified as a putative “chimera” if they have multiple inconsistent alignments with at least 10% of read length and 70% identity.

**Table S3. Properties of long reads from PacBio and ONT**. “PB” indicates PacBio. Reads were mapped to the assemblies using minimap2 to determine identity. We considered the average identity for each base, and if there were multiple alignments at a base, we used the one with the best score. We aligned PacBio reads to the hybrid assembly obtained from all PacBio reads. We aligned ONT reads to the hybrid assembly obtained from all ONT reads. N50 represents the length or identity for which half of the read bases are in reads of at least such length or identity.

**Table S4. Assembly runtimes in minutes**. All assemblies were run with dual 8-core Intel IvyBridge 2.6GHz, 256GB 1866MHz memory. Times include running times for Canu correction and read filtering.

**Table S5. Location and counts of proteins found uniquely in (a) ONT-based or (b) PacBio-based assembly for each sample**. Shown here is the comparison between assemblies using the globally optimal long read preparation strategy for each sequencing approach i.e. “Subsampled” for ONT+Illumina and “Basic” for PacBio+Illumina (as in Figure S4). Proteins from assemblies for each sample were clustered using Roary after annotation with Prokka. Contig order indicates size order in the relevant assembly (see Figure S4). The start of the greyed-out squares indicates the total number of contigs in the assembly.

**Table S6. Results of DNAdiff comparison between reference genomes (*E. coli* CFT073 and *K. pneumoniae* MGH78578 genomes) and hybrid assemblies with either PacBio or ONT**. Each row corresponds to a comparison, either between the reference and PacBio assembly, or between the reference and the ONT assembly, or between the two *de novo* hybrid assemblies. “Length difference” means the difference in total length of the two genomes. “Aligned bases (ref)” represents the number of bases from the first comparison genome that are aligned with the other genome in the comparison. In each comparison the ONT assembly is the one obtained using half of the long reads, while the PacBio assembly is obtained following long read correction.

**Table S7. Simulating the effect of increased level of ONT multiplexing on hybrid assembly**. Values represent numbers of contigs, either circular contigs, or any contig. Three simulations are presented, either with all reads, with half the reads, or with one third of the reads.

