## Supplementary figures and images for "Comparison of long-read sequencing technologies in the hybrid assembly of complex bacterial genomes"

### Figure S1

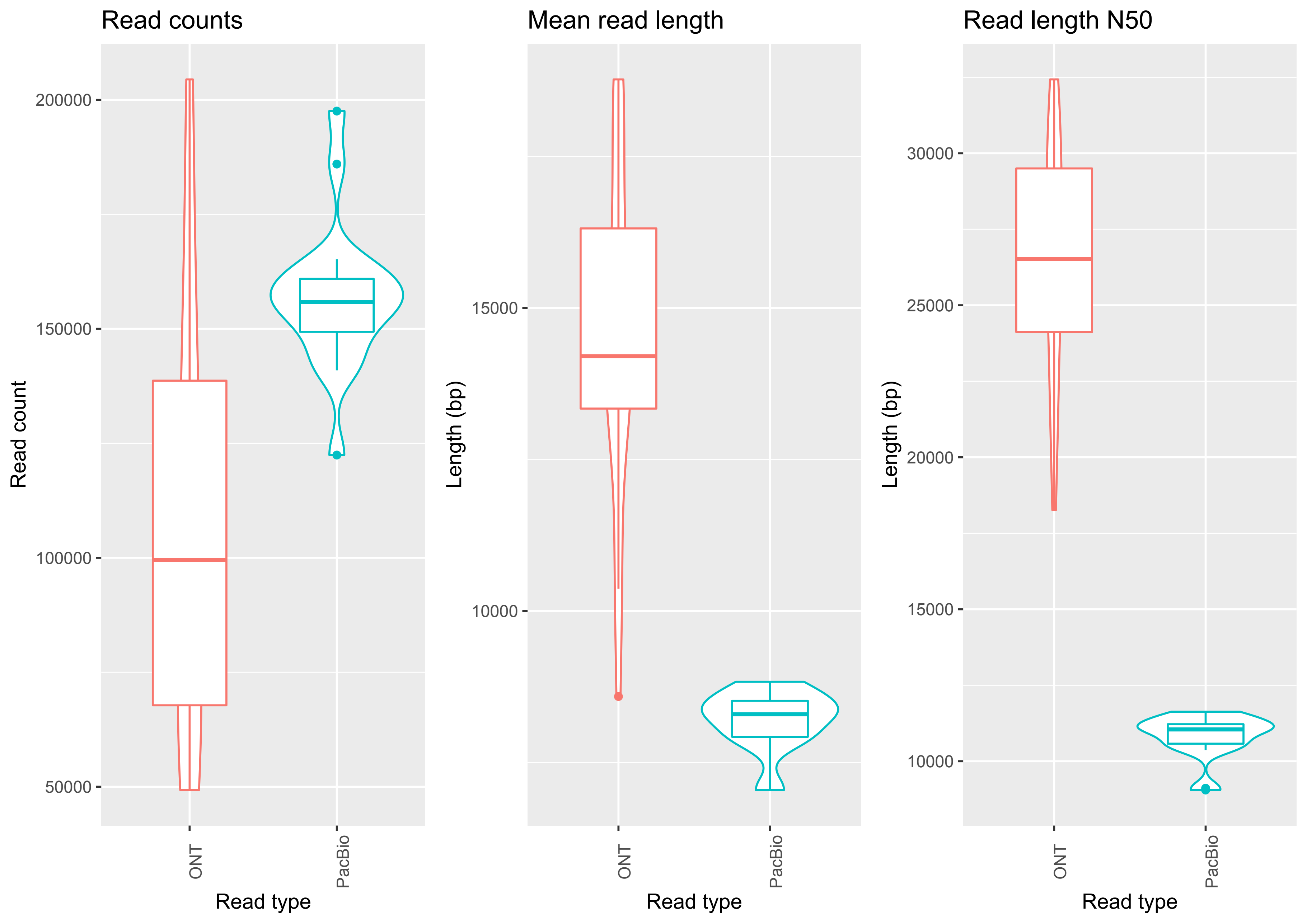

### Figure S2

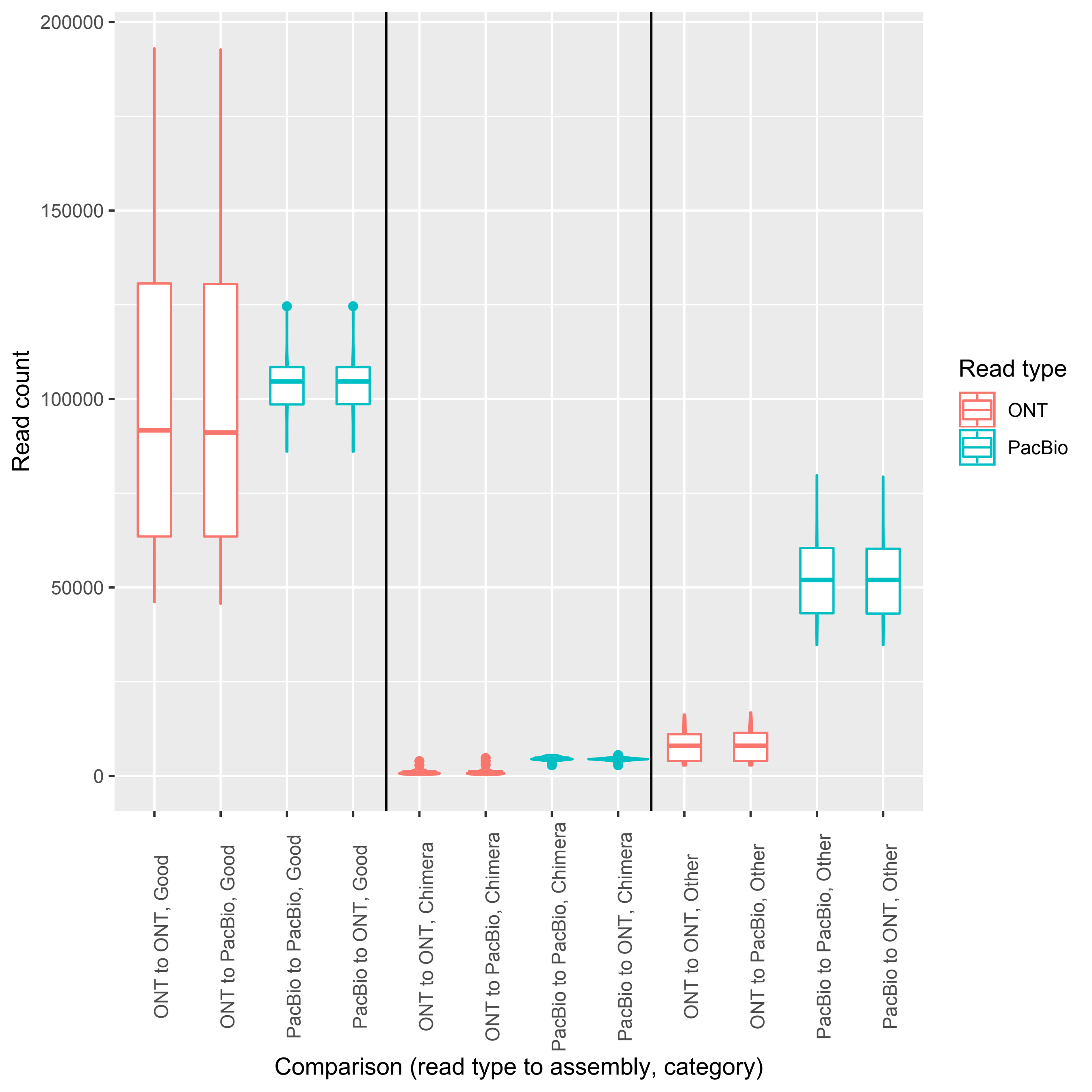

### Figure S3

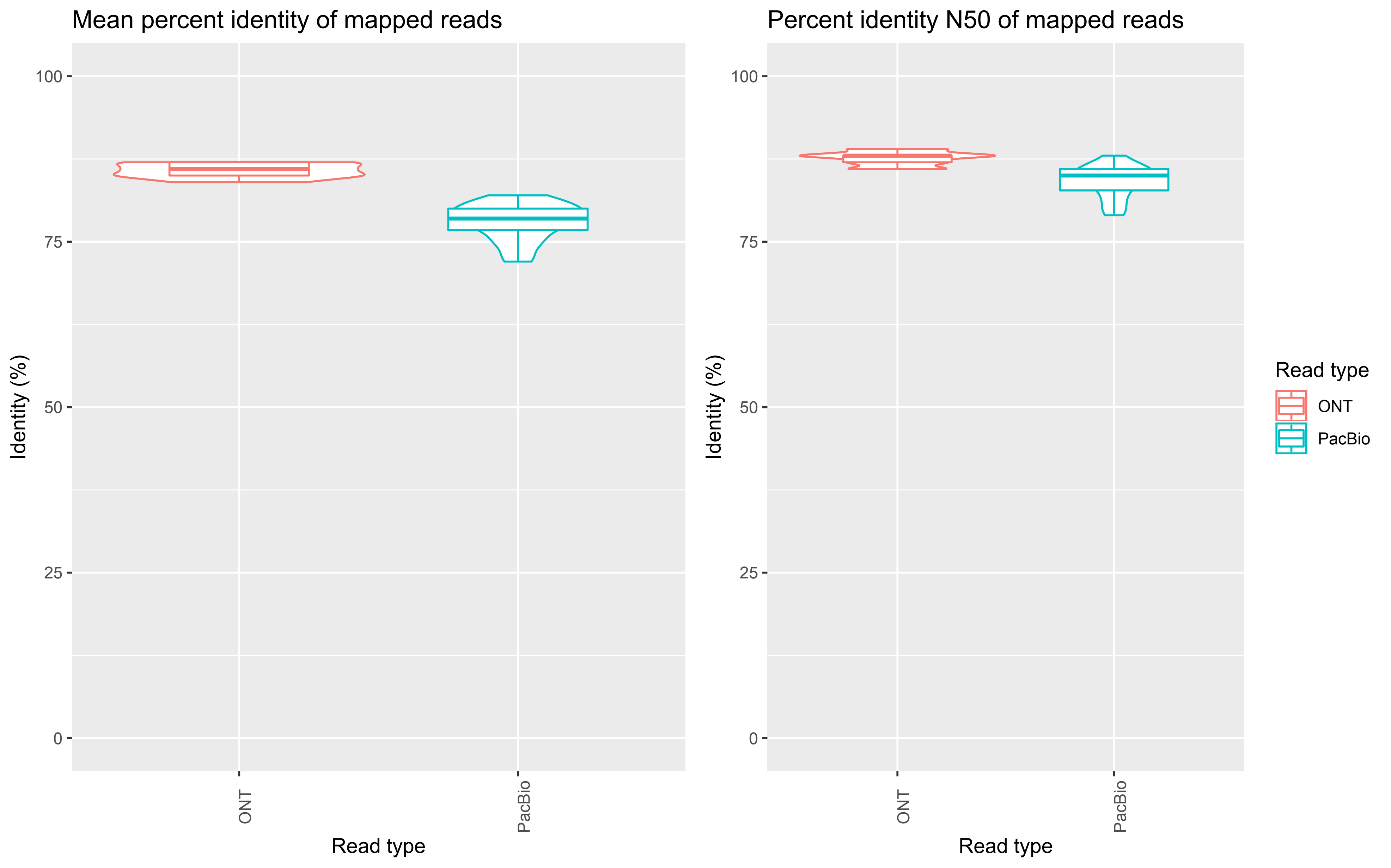

### Figure S4

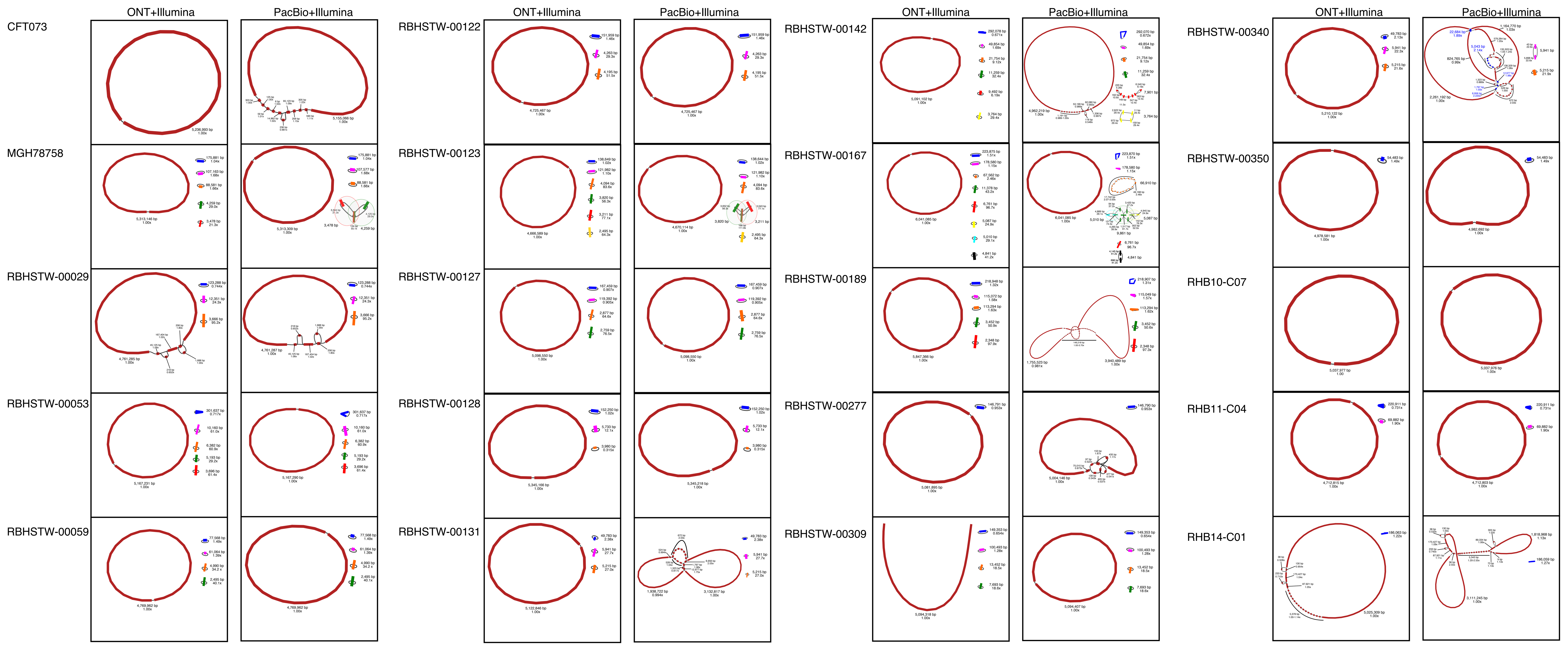

### Figure S5

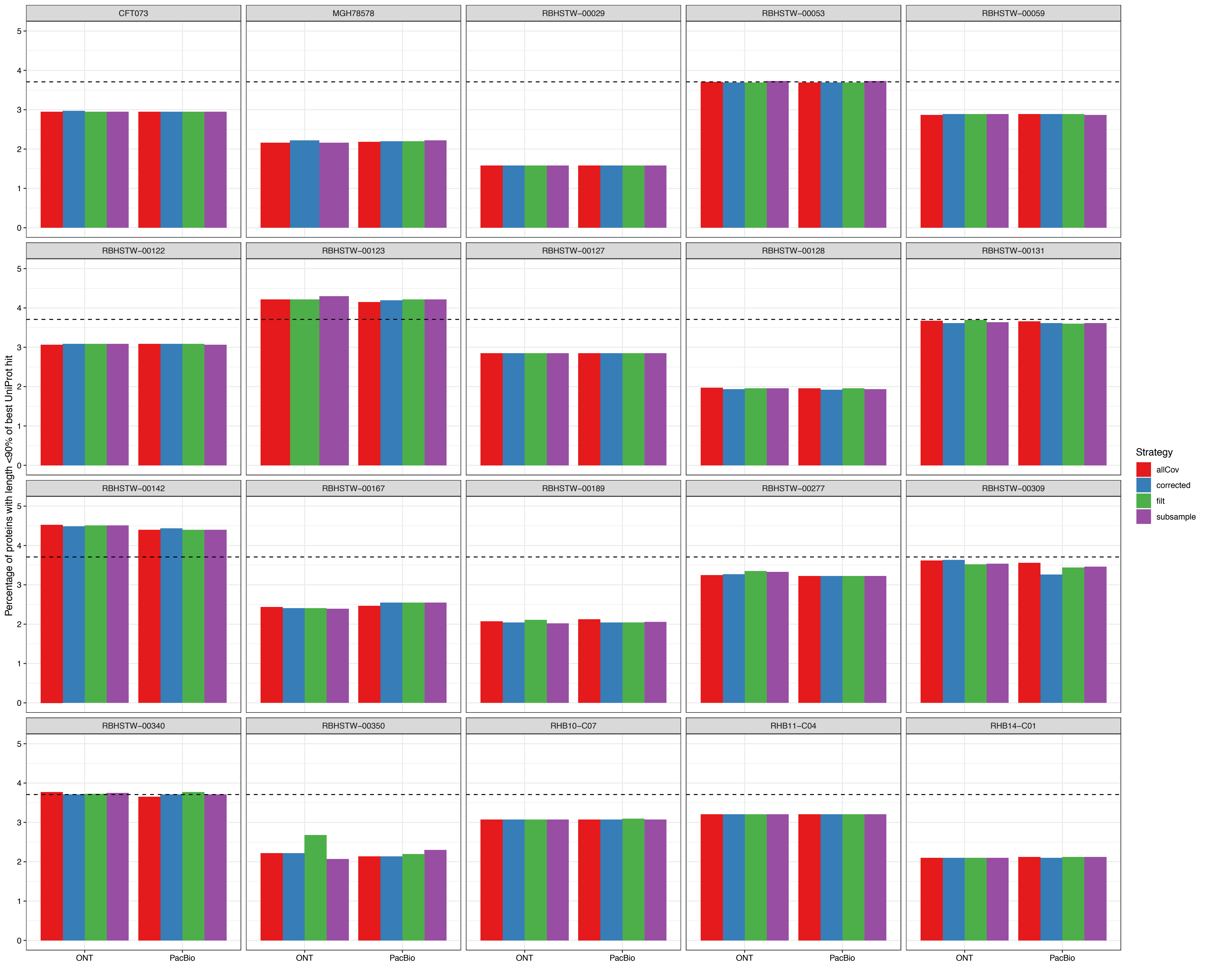

### Figure S6

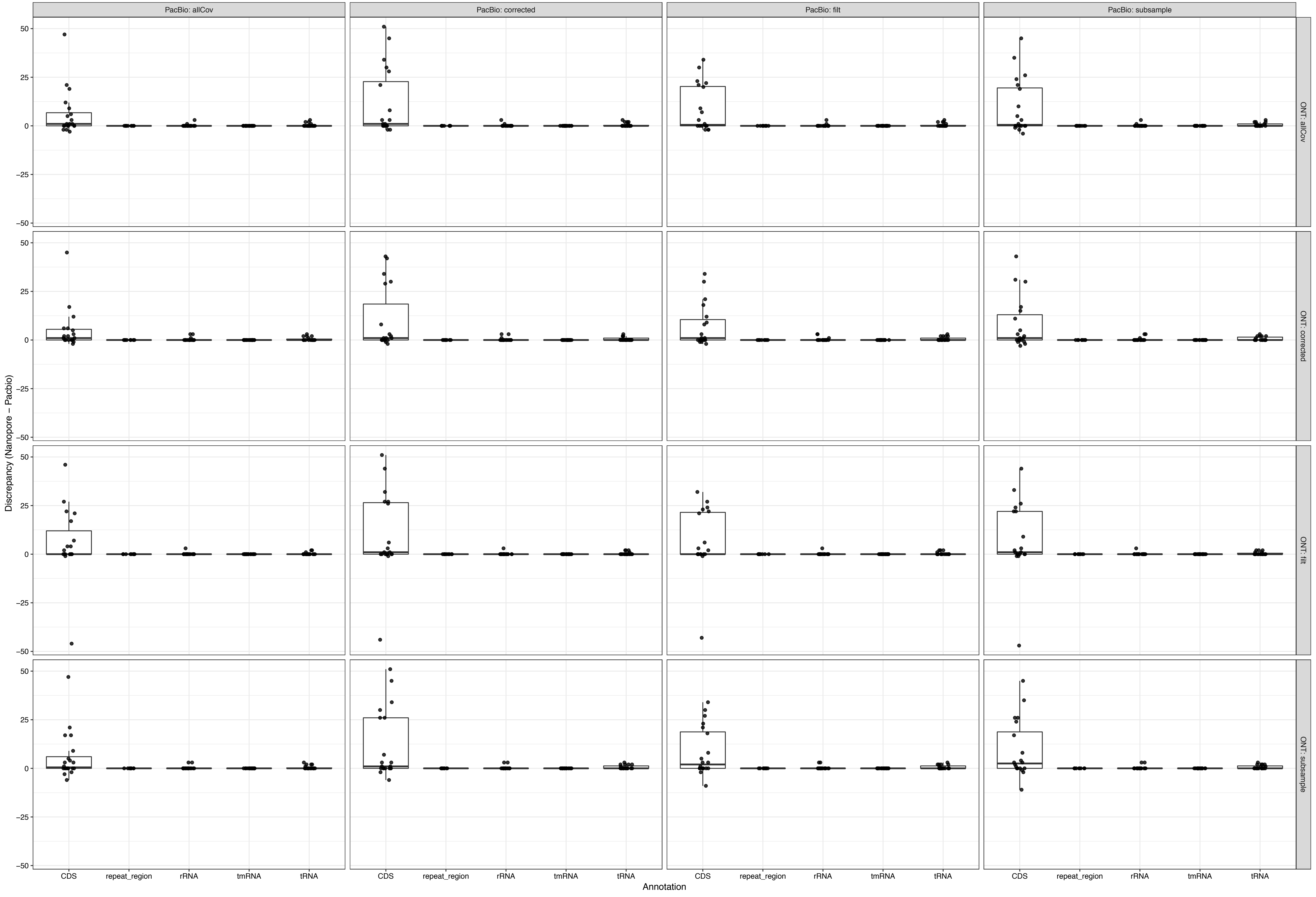
